# From impact metrics and open science to communicating research: Journalists’ awareness of academic controversies

**DOI:** 10.1101/2024.09.03.609638

**Authors:** Alice Fleerackers, Laura L. Moorhead, Juan Pablo Alperin, Michelle Riedlinger, Lauren A. Maggio

## Abstract

This study sheds light on how journalists respond to evolving debates within academia around topics including research integrity, improper use of metrics to measure research quality and impact, and the risks and benefits of the open science movement. Drawing on semi-structured interviews with 19 health and science journalists, we describe journalists’ awareness of these controversies and the ways in which that awareness, in turn, shapes the practices they use to select, verify, and communicate research. Our findings suggest that journalists’ perceptions of debates in scholarly communication vary widely, with some displaying a highly critical and nuanced understanding and others presenting a more limited awareness. Those with a more in-depth understanding report closely scrutinizing the research they report, carefully vetting the study design, methodology, and analyses. Those with a more limited awareness are more trusting of the peer review system as a quality control system and more willing to rely on researchers when determining what research to report on and how to vet and frame it. We discuss the benefits and risks of these varied perceptions and practices, highlighting the implications for the nature of the research media coverage that reaches the public.

---

The systems of scholarly communication and journalism are deeply intertwined, with changes in one influencing the values, norms, and practices of the other [e.g., 1,2]. Yet, the mechanisms underpinning these interconnections are not well understood, with a lack of evidence into how journalists make sense of, and adapt their work in response to, changes in scholarly communication [3]. To help fill this gap, we present a qualitative analysis of 19 interviews with health and science journalists examining their perceptions of key scholarly communication debates (e.g., on “impact” metrics, research integrity, open access [OA]) and how these perceptions, in turn, relate to the professional norms, practices, and criteria they use to select, verify, and communicate research. In doing so, we shed light on the consequences of the increasingly close coupling between science and journalism [4,5] for scientists, journalists, and the public.

## Close relationships and heavy reliance between science and the media

Decades of research have demonstrated the close and codependent relationships that can develop between science journalists and the researchers whose work they report on [5,6]. These studies show that science journalists rely heavily on interviews with researchers to help them identify new story ideas, vet the quality of studies, contextualize and translate the findings for public audiences, frame the implications, and more [7]. Journalists sometimes also rely on researchers for access to research papers that are behind paywalls [8,9]. Researchers, in turn, rely on journalists to share new findings to the public, gain societal legitimacy, and win funding—a reliance that may be increasing as scientists and scientific institutions face growing pressures to demonstrate a wider societal “impact” of their work [10,11].

Journalism and academia share many commonalities—including a shared commitment to independent, rigorous investigation in pursuit of “truth”—but they are also distinct in important ways, including differing epistemologies, ethical frameworks, and standards of practice (Bartleman et al., 2024). As such, although the close relationships between journalists and scientists may enable them to work together more efficiently [12], they also come with risks. In particular, scientists may prioritize the norms, values, ethics, and practices—or *logics*—of journalism over those of science, potentially resulting in a loss of scientific autonomy with respect to how research is conducted [13] and communicated [11,14,15]. For journalists, internalizing the logics of science may result in media coverage that prioritizes the interests and perspectives of researchers over those of the public. Moreover, journalists’ heavy reliance on scientists could prevent them from reporting what is relevant to society (i.e., rather than what is relevant for particular scientists or scientific institutions). It could also compromise their ability to provide a balanced portrait of reality that gives space for competing interpretations or dissenting perspectives or act as watchdogs of the powerful (e.g., scientists, funders, pharmaceutical companies) [16].

Understanding the potential risks and benefits of journalists’ proximity to scientists is especially important given growing concerns about research integrity, efforts at research assessment reform away from the use of “impact” metrics, and broader changes in scholarly communication resulting from the open science (OS) movement [17]. For instance, a nuanced understanding of problematic aspects of science (e.g., research fraud, retractions, improper use of metrics) is essential if journalists are to effectively perform their watchdog role—i.e., to sound an alarm when the actions of scientists put the public interest at risk. Yet, a more incomplete understanding of these issues could result in media coverage that inaccurately portrays research infractions of evidence that science is “broken,” damaging public trust [18,19]. Similarly, journalists’ awareness of OS practices—such as the use of preprints, OA papers, or open data sets—may facilitate their work by enabling them to use research knowledge that would otherwise be inaccessible [20–22]. Yet, a more superficial understanding of these practices, or one that prioritizes the interests of science over those of the public, could result in problematic media coverage, such as stories conflating the OA movement with predatory publishing [23,24] or stories that present preprints as if they were peer reviewed research [25–27].

Despite these potential risks and benefits, we lack research that has examined the *scientization* of journalism—that is, “the influence that researchers exert on the media as they vie for their research (or opinions) to be featured in lieu of other coverage” [3]. This exploratory study contributes to filling this gap by documenting journalists’ awareness of key controversies within academia, and how this understanding, in turn, shapes their practices for reporting on science.

## Evolving debates within scholarly communication

In recent years, changes in the scholarly communication ecosystem have sparked debates about interrelated concerns regarding research integrity, challenges of assessing research quality, and adverse effects of impact metrics, as well as on the potential role of OS to either address or exacerbate these concerns. These concerns, and the debates around them, are complex, involving actors at every level of scientific systems, especially researchers and administrators at academic institutions.

Underscoring them all is a “publish-or-perish” culture that exerts pressure on academics to produce as many research articles as possible and ensure they are published in prestigious and “high impact” journals [28,29]. This pressure has led to a hyper-competitive academic environment that relies heavily on mechanisms for tracking and assessing research and researchers for resource allocation (e.g., career progression, departmental funding, etc.) [30]. The effects of these practices are system-wide and form part of a larger set of changes that some have described as the neoliberalization of the university [31,32]. Addressing the full scope of these discussions and concerns is beyond the scope of this literature review, but we touch on the more direct concerns in the following paragraphs, as they pertain to how scholars understand and talk about their work with each other and, potentially, with the journalists they engage with.

On a pragmatic level, the growing pressures to publish have fueled multiple concerns about research integrity. Early among these was the so-called “replication crisis” stemming from the high volume of studies whose findings could not be verified, either due to lack of methodological detail or accessible data or because results were not robust enough [33]. More recently, the rise of a pay-to-publish model fueled the rise of so-called “predatory journals” that charge authors but do not offer a legitimate peer review process or follow other editorial best practices [34,35]. Today, one of the most pressing concerns is the emergence of “paper mills” that sell co-authorships in real, or sometimes machine-written, articles that authors can add to their CVs [36,37]. These specific concerns and the resulting debates have led to multiple responses from the academic community, including greater calls for research assessment reform [38], close scrutiny of the peer review system [39], and efforts towards greater openness and transparency [40].

One place to enact such changes in research assessment is in review, promotion, and tenure (RPT) processes, which often purport to reward many aspects of academic work but are known to place emphasis on the number of publications [41] and the conflated concepts of prestige, quality, and impact [42]. Proposed alternatives include the use of narrative CVs (i.e., allow those being assessed to describe their career trajectory) [43], development of values-based indicators [44], and other assessment frameworks that place emphasis on the quality over quantity of research outputs [45].

Generally, calls for research assessment reform have focused on pushing back on the simplistic use of citation-based metrics, especially the Journal Impact Factor (JIF) [38,46]. While there is broad consensus on the limitations and adverse effects of the JIF [47], its use for research assessment remains widespread [28], in part because of the labour of assessing the quality of a growing number of publications. Eve and Priego [48] argue that relying on these indicators is driven by a need to conserve reading labour: the metrics let scholars decide what, in the vast body of literature, is worth reading, and who among their peers is worth rewarding.

Peer review is another way in which the scholarly community has traditionally delineated between problematic and reliable or “authoritative” scholarship [49]. Yet, like journal metrics, the effectiveness of peer review itself has been subject to debates in recent years. These debates largely centre on the (in)effectiveness and (in)appropriateness of peer review for weeding out problematic research, but also include other considerations such as biases in the review system and their impact on the diversity of the scholarly record; the burden of peer review on already strained academics; and the lack of transparency involved [39,50–52].

Ongoing discussions about the role of OS in scholarly communication have brought another layer of complexity to these debates about research integrity, impact, and assessment. Proponents of OS argue that practices such as sharing research data and protocols or publishing open peer review reports can bring transparency to the research and publication process and, in turn, improve the integrity of the scholarly record [53], shift the scholarly community’s reward system toward valuing a more diverse range of research outputs [54], or more broadly help to realign incentives as part of the push for research assessment reform [55]. Advocates also see OS as a mechanism for making scholarly communication more equitable and inclusive, as openness can allow less well-resourced scholars to access and, for some forms of OS, contribute to research knowledge [56,57]. Yet others have raised concerns that OS may further exacerbate existing problems in academia. For example, the high cost of article processing charges required to publish OA in some journals could compound existing inequities in whose research is published and rewarded [58]. Moreover, some fear that the growing use of un-peer-reviewed preprints in scholarly communication’ as was seen during the COVID-19 pandemic, could contribute to the spread flawed science, misinformation, and even conspiracy theories [59,60].

In sum, in reporting on research, journalists must navigate a contentious and rapidly evolving scholarly communication landscape and an ongoing questioning of how research should be vetted, valued, and rewarded. Due to their close relationships with scientists, it is likely that these challenging debates influence how journalists think about and cover research; yet the nature of this influence is not yet well understood. To address this important gap, we interviewed 19 health and science journalists about their reporting practices and perceptions of research. Our results provide some of the first evidence into how journalists have internalized key controversies in scholarly communication and how this internalization shapes ways in which they report on research.

## Methods

Guided by a constructivist paradigm, we conducted an interview study using qualitative description [61], meaning that we aimed to gather data that describe the “why, how, and what questions about human behavior, motives, views, and barriers” [62]. We selected this methodology as it is well suited for use when interviewing individuals with direct experience of the studied phenomenon.

This study is a component of a larger research project in which we conducted interviews with journalists and scientists. Given our research aim, this current investigation focuses only on the journalists’ interviews. For further details of the larger project see Fleerackers et al. [63] and Moorhead et al. [12]. The Simon Fraser University Research Ethics Board (# 30000244) and the San Francisco State Institutional Review Board (#2021175) exempted the project from further review.

### Sample

Eligible participants for this study worked for one of seven news outlets that included a mixture of traditional, legacy news organizations (i.e., *The Guardian, New York Times*); historically print-only science magazines (*Popular Science, Wired*); digital-native health sites (*News Medical, MedPage Today*); and a science blog (*IFLScience*). We selected these news outlets based on their focus on science and health news and frequent coverage of academic research. Their diversity (in formats, publishing models, audiences) was also desirable because it was more representative of today’s digital media ecosystem than a sample of traditional, legacy news outlets. Journalists from these outlets were eligible to participate if they had published a story between March 1 and April 30, 2021, that included a mention of research. Mentions of research were identified through the outlet’s RSS feed or via the Twitter timeline of the official account that posted a link to every story. We have made the scripts used for this process openly available [64].

### Data Collection

Between July and November 2021, a research assistant with experience in journalism emailed eligible journalists to participate using publicly available email addresses. Nineteen journalists were interviewed (see Fleerackers et al. [63] for participant characteristics). Journalists consented to participate in the study via a consent form sent before the interviews were conducted.

The research assistant conducted and recorded semi-structured interviews with the journalists via Zoom. In the interviews, participants were asked to describe their professional experience reporting on research, including how they engage with scientists. The interview guide is available online [65].

Interviews were on average 35 minutes (range: 10-47 minutes). A third-party transcriptions service transcribed all transcripts, which were then de-identified by LLM prior to analysis.

### Data Analysis

Analysis of the data began with AF identifying and highlighting sections of the transcripts broadly applicable to the participants’ descriptions of their engagement with scientific debates and practices (e.g., searching for studies via academic databases, evaluating research quality based on methodological criteria, placing trust in peer review, valuing citation counts or journal reputation, supporting the OS, etc.). All authors were provided access to the complete, highlighted transcripts and tasked with familiarizing themselves with the overall content and conducting open coding of elements that might relate to journalists’ understanding and adoption of scholarly debates and norms. The authors then met virtually to discuss and co-create a working codebook, which was subsequently refined by AF after additional readings of the transcripts to include draft definitions and examples of codes (i.e., codebook thematic analysis) [66]. The authors then independently applied the codebook to the transcripts, while staying open to the identification of additional codes, and again met to discuss areas requiring revision. Informed by these conversations, familiarity with the data, and literature on journalism and scholarly communication, AF iteratively refined and applied the newly organized codes to transcripts, and, eventually, solidified the codebook.

To explore potential patterns in how different types of journalists understood academic debates and how this affected their work, AF did a final round of coding using the qualitative analysis software NVivo 12 [67], which allows researchers to easily assess whether codes and themes are expressed by all participants (i.e., cases) or by only a subset with particular demographic characteristics via its crosstabs query functionality. Examining potential differences among types of journalists was important as some of the participants had professional training or education in science (and thus would be theoretically more likely to be aware of controversies in academia), while others did not. Similarly, some worked as specialized science journalists at science-focused outlets, whereas others reported on other beats (e.g., lifestyle, culture) or worked for other types of outlets (e.g., general news outlets). The specialized journalists could potentially be more sensitized to scholarly debates and norms than those with less experience reading academic research or interacting with scientists. While our qualitative design did not allow us to quantitatively compare different groups of journalists, we assessed whether evidence of each theme was present among each type of journalist. Specifically, we looked for the presence of each theme among journalists of different educational backgrounds (i.e., in journalism, science, both, or some other field), levels of education (i.e., bachelor’s, master’s, doctoral, unknown), years of experience (i.e., 0–4, 5–9, 10+), roles (i.e., staff reporter, freelancer, staff editor), and beats (i.e., science/health vs other). We also assessed whether themes were expressed by journalists working at outlets of different types (i.e., traditional/legacy journalism vs alternative/peripheral media)[68] and different topic specializations (i.e., health/science vs other). The final themes (described in the Results and Discussion section) were expressed by journalists within each of these subgroups.

### Reflexivity

Our research team included individuals with a variety of backgrounds and experiences as researchers (all authors), journalists (AF and LLM), and communication professionals (AF and MR) who have worked in North American contexts. JPA’s research focuses on scholarly communication, while LAM studies the prevalence of irresponsible research practices by scientists. Additionally, all members of the research team shared a background and interest in OS and the responsible use of research metrics. Taken together, the research team’s interests and expertise likely influenced the study aims, analysis, and conclusions.

## Results and Discussion

Our results suggest that journalists are generally aware of, and influenced by, debates about issues such as research integrity, impact metrics, and OS that are taking place within academia. Yet, journalists vary widely in terms of how closely they are enmeshed with these debates, with some presenting a highly critical and nuanced understanding and others presenting a more limited awareness. These different levels of awareness contribute to different approaches to reporting on research. We report these varied perspectives and practices below and discuss their implications for the nature of science media coverage. To provide context for our findings we provide direct quotes from participants across our sample. Participants are identified by their participant number (e.g., J11 is journalist participant 11).

### Critical Awareness of Academic Debates

Interviews with journalists revealed that some had a deep awareness of ongoing debates and discourses taking place within the scholarly community—an awareness that sometimes influenced how they selected, verified, and communicated research. For example, journalists expressed a nuanced understanding of the limitations of peer review, including its slow, potentially biased, and often imperfect effectiveness as a quality control mechanism [39,69]. Others referenced related challenges to research integrity, such as the ongoing replication crisis, the rise of predatory journals, the pressure to publish and its implications for research quality, and the problematic nature of publication bias. J11, for instance, noted that “there are certain—what’s the word—certain pressures, you know, to sort of publish or perish and that people might be kind of cranking out a lot of articles.” This awareness of ongoing challenges to the integrity of science led some journalists to report findings with additional scrutiny.

This was the case for J1, for example, whose knowledge of the replication crisis made them skeptical of using older research if “there’s no follow-up” or newer studies that have come to similar conclusions.

Perhaps because of their awareness of research integrity issues, journalists described relying on scientific methods for ensuring the research they reported was accurate, unbiased, and trustworthy. This included, above all, a consideration of the study design and methodology. Although several journalists noted that they lacked the expertise to effectively verify complex statistical analyses [70], they still attempted to scrutinize the quality of the research before deciding to report on it, something that is notoriously difficult to define and assess, even from within the scientific community [71]. Common strategies included critically reading the paper and investigating the nature and size of the sample; human subjects were valued above mice, and larger samples were generally seen as more trustworthy than smaller ones. (One journalist, however, noted that small samples are appropriate in qualitative studies, demonstrating a more nuanced understanding of research methods and study design.) Meta-analyses and randomized controlled trials (RCTs) were preferred over other types of studies, which aligns with how researchers conceptualize the hierarchy of available evidence [72,73]. Journalists looked for causality, in ways similar to scientists, asking questions such as, “Is there an implicit bias in the study? Are the conclusions valid? Is it really a causation, for example, or just a correlation?” [J12] and noting whether “their methodologies leave this open to be, like, a correlation, not necessarily a causation” [J15]. Finally, some journalists saw research as more credible when researchers noted study limitations and did not have any clear conflicts of interest, aligning with the consensus building in the scientific community that transparent reporting is an indicator of research integrity [74]. A few journalists went even further, running their own statistical tests on publicly available datasets to provide new insights. These journalists made comments such as, “I was very much playing the scientist, almost, in that I was the one writing, ‘What does the data mean?’” [J2].

Journalists’ critical understanding of academic debates extended beyond issues of research quality. For example, some journalists noted the increasing need for universities to establish legitimacy and gain public support through self-interested science public relations efforts, echoing similar findings to those described by Weingart [11]. This was reflected in comments such as “[Research papers] come to me either directly if I’ve worked with them before or through their press release offices, right, which are pumping out research all the time in order to garner interest” [J4]. Others, such as J12, expressed awareness of the importance of establishing priority as a scientist and avoiding getting “scooped.” This knowledge, in turn, informed how and when they reported on scientists’ research:

> …you have to cover [a] preprint as soon as possible…Give the researchers that put the preprint also another, how to say it, another stamp of—they were the first—that their research was the first, because sometimes, that can be problematic in scientific circles [J12].

Even more common were discussions about the public’s right to access research—a core pillar of the OS and OA movements that have been gaining momentum in academia over the past two decades [57]. These reflections on the public’s right to research knowledge were often grounded in journalists’ own frustrations in accessing the literature, as evidenced by J19, who stated that “it’s so frustrating when you finally find the paper that you want to read and it’s behind a paywall, particularly when it’s publicly funded research, you know?”

Yet journalists also discussed OA in other contexts, such as when reflecting on the rise of preprints during the COVID-19 pandemic. Journalists expressed an awareness of the benefits that scientists find in preprints, as both a rapid-sharing mechanism that avoids the slow peer review and publication process, and as a way to circumvent paywalls from costly subscription journals [75]. As J16 explained,

> [Preprints are] the way that scientists talk quickly to each other, and especially as people realize that they don’t have easy access to the big journals because they cost a lot of money and they’re behind paywalls, that this becomes much more of a way for scientists to talk to each other and to a potential audience for that work.

### Limits in understanding of academic debates

Journalists’ in-depth understanding of academic debates was not universal, however. For example, alongside journalists who were skeptical of peer review were journalists who saw it as the “gold standard” [J13] to guarantee that findings were trustworthy and credible enough to report on. These journalists made comments such as, “with the peer reviewed piece, you’re really saying ‘This is a discovery or something that is fairly well-vetted and legitimate’” [J18]. Scholars have previously noted that journalists use peer review as a stamp of approval, one which moves the findings from uncertain evidence to confirmed facts [6,76,77]. This view of peer review aligns with that of many researchers [78] despite criticisms that peer review is not a reliable quality control system and can, at times, introduce biases [39,50].

Similarly, while some journalists were aware of the pressures researchers faced to demonstrate “impact,” others uncritically accepted and relied on measures of impact used by many scholars. Specifically, journalists described selecting studies to report based on indicators such as the number of citations an article receives, the impact factor of the journal it is published in, and the reputation of the journal, the authors, and their institutions [29,79]. This reliance on proxies for research quality and impact can be seen in comments such as “you can do stuff to verify how, the impact factor of a journal or how legitimate the research is” [J9] and “that’s why they’re in *Science* and *Nature*, ’cause they’re big stories, they’re important” [J5]. As J10 explained, over time, these indicators formed an internalized framework that journalists could fall back on without having to critically scrutinize study findings:

> We’ve got, like, a fairly good understanding of we know the journals that you almost don’t have to question too much, you know, like, the *Nature* journals, *Science*, *PLOS ONE*, all the *PLOS* ones, you kind of realize, like, these are fairly well-respected, so you don’t have to question them too much.

The focus on reputational assessment strategies aligns with findings of Badenschier and Wormer [80] and Rosen et al. [81], who similarly identified “scientific relevance” as a value shaping science journalists’ selection decisions. Deferring decisions about the impact or quality of scientific outputs to recognized scientific community mechanisms potentially saves time for journalists, but our findings reveal how doing so can enable problematic aspects of academia to seep into journalism, such as an overreliance on citation-based metrics of “impact” and journal reputation [48]. Citation indicators such as the JIF are known to be poor measures of the quality of individual articles and to be biased against journals in the social sciences and humanities, as well as those from the Global South [47,82]. The use of these metrics by journalists could serve to further perpetuate their problematic use within academia.

Similarly, while some journalists were aware that scientists faced pressures to promote their work, this did not translate to a critical relationship with press officers or the scientists whose work they promoted. Instead, journalists looked to journal and university press officers to understand the research, as reflected in comments such as:

> I couldn’t do my job without the public information officers. They’re great, and especially for some of the studies that are written in a language that even I find quite difficult to access, like nothing is better than a press release to just get your head straight about what the top-line findings of something are. [J18]

Similarly, none of the journalists in our sample mentioned the importance of maintaining independence from the scientists they interviewed, even though independence from sources is one of journalism’s core values [83] and a foundational ethical principle of science journalism [84]. Instead, journalists described their relationships to scientists as collaborative and trusting, as reflected in statements such as, “Obviously, I have my own sources, and if I have a source who’s in that topic area, I would reach out to them,” [J17] and, “I have sources that I rely on and that I think are trustworthy, and I go to them” [J7]. Journalists expected scientists to act as unbiased experts on the topic at hand, even while acknowledging their fallibility. As J7 noted, incorporating expert opinions “gives [the story] credibility that makes me feel better, even though, yes, they sometimes are not all that accurate” [J7].

As has been noted in previous research [6,63], journalists leaned heavily on these trusted sources to critique and vet other researchers’ work. Yet, journalists relied on scientists for more fundamental aspects of their reporting as well. Many asked experts to translate the research—to “give more insight and clear explanation to the article that we could provide by ourselves” [J1]. This was especially important for journalists without a background in science, such as J11:

> One thing is that, again, often kind of coming in as a liberal arts backgrounded person with not a huge understanding of something like computational fluid dynamics, that I would hope that they are a resource I can trust, you know, and, again, that gets into a tricky thing, because, you know, if I can’t tell that they’re wrong, you know, who’s going to tell me they’re wrong?

Journalists also used scientists when seeking out research to cover or incorporate in their stories, “relying on the expertise of people who research in that area to point me towards the like, external literature to begin with” [J14]. On a practical level, journalists also frequently used scientists to get access to (paywalled) research papers [9,85]. This was true even of participants working at major publications, such as the staff reporter at *The Guardian* who reported that, “A lot of the time, it’s really hard to find a PDF of the paper, and, like, scientists are brilliant. If you email them, they send you a PDF of the paper” [J19]. In this sense, scientists not only acted as collaborators in journalists’ work [12] but also as gatekeepers and agenda setters. Their actions and input helped shape what research got coverage, as well as how it was contextualized and framed [86]. Scientists, scientific institutions, and scientific journals were assumed to stand by the research findings they communicated with conviction.

Science journalists’ overly reliant relationships on expert sources have been discussed elsewhere [84,87] and align with the narrower definitions of scientization used in previous research—i.e., the increasing influence of scientific and expert sources on journalists’ work [3,88]. Our findings extend this previous research by providing a view into how and why journalists work with particular scientists and how they come to negotiate this intersection of seemingly disparate professions.

### Limitations

These findings must be considered in light of several limitations. First, the journalists we interviewed responded to the recruitment email, which suggests they likely had an existing interest in science relations and may thus have been more enmeshed in scholarly communication debates than others. They also represented a specific subset of journalists—those who reported on research at least occasionally, produced text-based stories, worked for online (rather than broadcast) media outlets, wrote in English, and were based in the Global North. The participants were also interviewed during the second year of the COVID-19 pandemic, when public interest in science was high [89] and concerns about research integrity were growing [90]. The practices and understanding of academic debates we identified may thus differ from those we might find among journalists working in other geographic, linguistic, professional, and temporal contexts. Relatedly, the qualitative nature of the study design means that it was not possible to systematically assess differences among journalists based on publication outlets, reporting interests, or other characteristics. Future research could test our framework with a larger sample of science journalists and a study design that would allow for comparisons across participant contexts and characteristics.

## Conclusion

In this study, we used in-depth interviews with journalists who report on science to further explore and understand the consequences of recent debates in scholarly communication and the close and codependent relationships that can develop between journalists and scientists that have been described by Franzen et al. [5], Weingart [11], Moorhead et al. [12], and others. In doing so, our analysis revealed how evolving academic debates not only influence how journalists perceive research studies (e.g., as valuable, impactful, trustworthy, or high-quality), they also shape their practices for reporting on them. Our findings present some of the first evidence of the impacts of the “scientization” of journalists described by MacGregor et al. [3], contributing to scholarly understanding of the interrelationships between science and journalism and shedding light on journalistic practices with important practical implications.

In some instances, journalists possessed an in-depth, critical understanding of academic debates about research integrity, which appeared to encourage a more critical approach to communicating science that could benefit their audiences. For example, some journalists expressed a deep awareness of the limitations of peer review and approached research studies with a skeptical eye. They decided which studies to report on by adopting practices used by many scholars, such as considering the study design and methodology and, in some cases, doing their own independent research (e.g., assessing the existing literature, collecting and analyzing their own data, and creating meaningful data visualizations).

At other times, however, journalists expressed a more superficial understanding of key challenges facing academia, which sometimes translated to problematic practices for reporting on research. For instance, some journalists described relying on academic citation metrics or markers of prestige as proxies for research quality—both practices that are themselves critiqued within academia [47]. None of the journalists we interviewed appeared to be concerned about maintaining independence from the scientists they were covering in their work, and most also trusted the information they received from university and journal press officers. This uncritical reliance on scientists and press officers was true even among journalists who acknowledged the potential fallibility of these sources and described the pressures facing scientists and scientific institutions to demonstrate the societal “impact” of their work. That is, at times, journalists’ reliance on scientific experts appeared to compromise their ability to serve the public interest and retain independence and autonomy from their sources—two commonly cited journalistic values [83,91].

Finally, journalists—as seen in the quotation “Who’s going to tell me they’re wrong?”—often recognized their limits in understanding and ability to vet research. Yet, their efforts to verify or triangulate research sometimes moved them closer to academia’s controversies and insular practices (e.g., press officers acting as knowledge brokers, scholars hesitant to speak on the work of peers, early-stage researchers desiring press coverage for their projects). Moving forward, there is a need to explore how journalists might better understand and document research integrity independently, without an overreliance on academia’s potentially limited and biased networks. Relatedly, our results suggest that advocates of OS have allies in science journalism, as journalists, too, appear to believe that research knowledge should be public knowledge. It is unclear, however, whether journalists align themselves with other OS goals, such as advancing transparency and improving research integrity, or make use of the increasing number of datasets, protocols, and other research outputs that are openly available [21]. As such, researchers may wish to explore how these open research outputs shape journalists’ practices for selecting, verifying, and communicating research to the public, as well as what new or additional tools and resources could be developed to support their work.

## Acknowledgements

We would like to acknowledge Mr. Asura Enkhbayar for his support in collecting the news stories which were used to identify potential participants and Ms. Kaylee Fagan Williams for her assistance in conducting the semi-structured interviews.

